# The Toxin Antitoxin MazEF Drives *Staphylococcus aureus* Chronic Infection

**DOI:** 10.1101/687434

**Authors:** Dongzhu Ma, Jonathan B. Mandell, Niles P. Donegan, Ambrose L. Cheung, Wanyan Ma, Scott Rothenberger, Robert M. Q. Shanks, Anthony R. Richardson, Kenneth L. Urish

## Abstract

*Staphylococcus aureus* is the major organism responsible for surgical implant infections. Antimicrobial treatment of these infections often fails leading to expensive surgical intervention and increased risk of mortality to the patient. The challenge in treating these infections is associated with the high tolerance of *S. aureus* biofilm to antibiotics. MazEF, a toxin-antitoxin system, is thought to be an important regulator of this phenotype, but its physiological function in *S. aureus* is controversial. Here, we examined the role of MazEF in developing chronic infections by comparing growth and antibiotic tolerance phenotypes in three *S. aureus* strains to their corresponding strains with disruption of *mazF* expression. Strains lacking *mazF* production showed increased biofilm growth, and decreased biofilm antibiotic tolerance. Deletion of *icaADBC* in the *mazF*::tn background suppressed the growth phenotype observed with *mazF*-disrupted strains, suggesting the phenotype was *ica*-dependent. We confirmed these phenotypes in our murine animal model. Loss of *mazF* resulted in increased bacterial burden and decreased survival rate compared to its wild-type strain demonstrating that loss of the *mazF* gene caused an increase in *S. aureus* virulence. Although lack of *mazF* gene expression increased *S. aureus* virulence, it was more susceptible to antibiotics *in vivo*. Combined, the ability of *mazF* to inhibit biofilm formation and promote biofilm antibiotic tolerance plays a critical role in transitioning from an acute to chronic infection that is difficult to eradicate with antibiotics alone.

**Importance:** Surgical infections are one of the most common types of infections obtained in a hospital. *Staphylococcus aureus* is the most common pathogen associated with this infection. These infections are resilient and difficult to eradicate as the bacteria form a biofilm, a community of bacteria held together by an extracellular matrix. Compared to bacteria floating in liquid, bacteria in a biofilm are more resistant to antibiotics. The mechanism behind how bacteria develop this resistance and establish a chronic infection is unknown. We demonstrate that *mazEF*, a toxin-antitoxin gene, inhibits biofilm formation and promotes biofilm antibiotic tolerance which allows *S. aureus* to transition from an acute to chronic infection that cannot be eradicated with antibiotics but is less virulent. This gene not only makes the bacteria more tolerant to antibiotics but makes the bacteria more tolerant to the host.

## Introduction

*Staphylococcus aureus* is a gram-positive pathogen associated with a variety of disease processes from self-limited abscesses to life-threatening sepsis. These episodes are typically acute, and resolve over a limited time-period with either minimal or high morbidity and mortality (1). An exception includes *S. aureus* related surgical infection, especially those associated with medical devices. Surgical site infection is one of the most common health-care associated infections (2). Unlike the majority of *S. aureus* infections, these infections can be chronic, indolent, and challenging to treat.

Periprosthetic joint infection illustrates this challenge. Total knee arthroplasty is a common surgical procedure, and the most common reason for failure is infection, termed periprosthetic joint infection (3, 4). *S. aureus* periprosthetic joint infection can be culture negative for prolonged periods (5, 6), has high failure rates above 50% once treatment is initiated (5), and a 5-year mortality of 20% (7–9), higher than many common cancers (10). Similar to other surgical implant associated infections, the challenge in treating this disease involves the ability of *S. aureus* to develop a chronic biofilm associated infection tolerant to antibiotics (11, 12).

In gram-positive bacteria, the mechanisms behind biofilm antibiotic tolerance and the ability to form chronic infections are poorly understood. It is suspected that toxin-antitoxin systems play an important role in these processes. Toxin-antitoxin (TA) systems encode a stable toxin protein capable of interfering with vital cellular processes and a labile antitoxin that counteracts the toxin (13–15). In *S. aureus*, the most well studied of these is the *mazEF* module where *mazF* is a stable toxin that cleaves specific mRNA, and *mazE* is an unstable antitoxin that inhibits *mazF* (16). In gram negative and acid-fast species, this system has been associated with antibiotic tolerance (17) and virulence (18). In *S. aureus*, the *mazEF* phenotype is controversial and its physiologic function in the disease process is unknown

The objective of this study was to identify a phenotype associated with *mazEF* in the *S. aureus* disease process. We hypothesized that toxin-antitoxin systems like *mazEF* contribute to the ability to establish chronic infections and antibiotic tolerant biofilms. Disruption of *mazF* expression in three different *S. aureus* strains, resulted in increased biofilm formation and a loss of antibiotic tolerance as compared to their wild-type strains on surgical implant material. In planktonic culture, when *mazF* disruption did alter growth, this was associated with antibiotic tolerance. In our animal model, the absence of *mazF* resulted in a more acute, pathogenic infection that was more difficult to treat with antibiotics. These phenotypes demonstrated that *mazF* expression resulted in lower growth and metabolic activity from decreased biofilm formation that allowed a transition from an acute to chronic biofilm infection and increased antibiotic tolerance.

## Results

### Disruption of *mazF* is associated with increased biofilm formation on surgical implant material

Toxin antitoxin systems are associated with bacteria growth arrest (19–21). We hypothesized that the lack of *mazF* would result in increased biofilm formation from preventing growth inhibition. Mature *S. aureus* (USA300 JE2) biofilm was cultured on titanium rods and quantitative culture was performed to asses biofilm mass. Disruption of *mazF* had increased biofilm mass as compared to parental strains (Fig. 1). We observed similar results on two additional methicillin sensitive *S. aureus* (MSSA) strains deleted for *mazF*, Newman (22) and SH1000 (23) (Fig. S1). These experiments were repeated, and biofilm was cultured on polystyrene, and quantified with crystal violet assay. A loss of *mazF* expression again resulted in increased biofilm mass on fibrinogen-coated wells in all three strains as compared to wild type (Fig. S2). To confirm the observed phenotype of *mazF* in *S. aureus*, we restored *mazF* expression *in trans* and observed a decrease in biofilm formation (Fig. S3).

**Figure 1.**
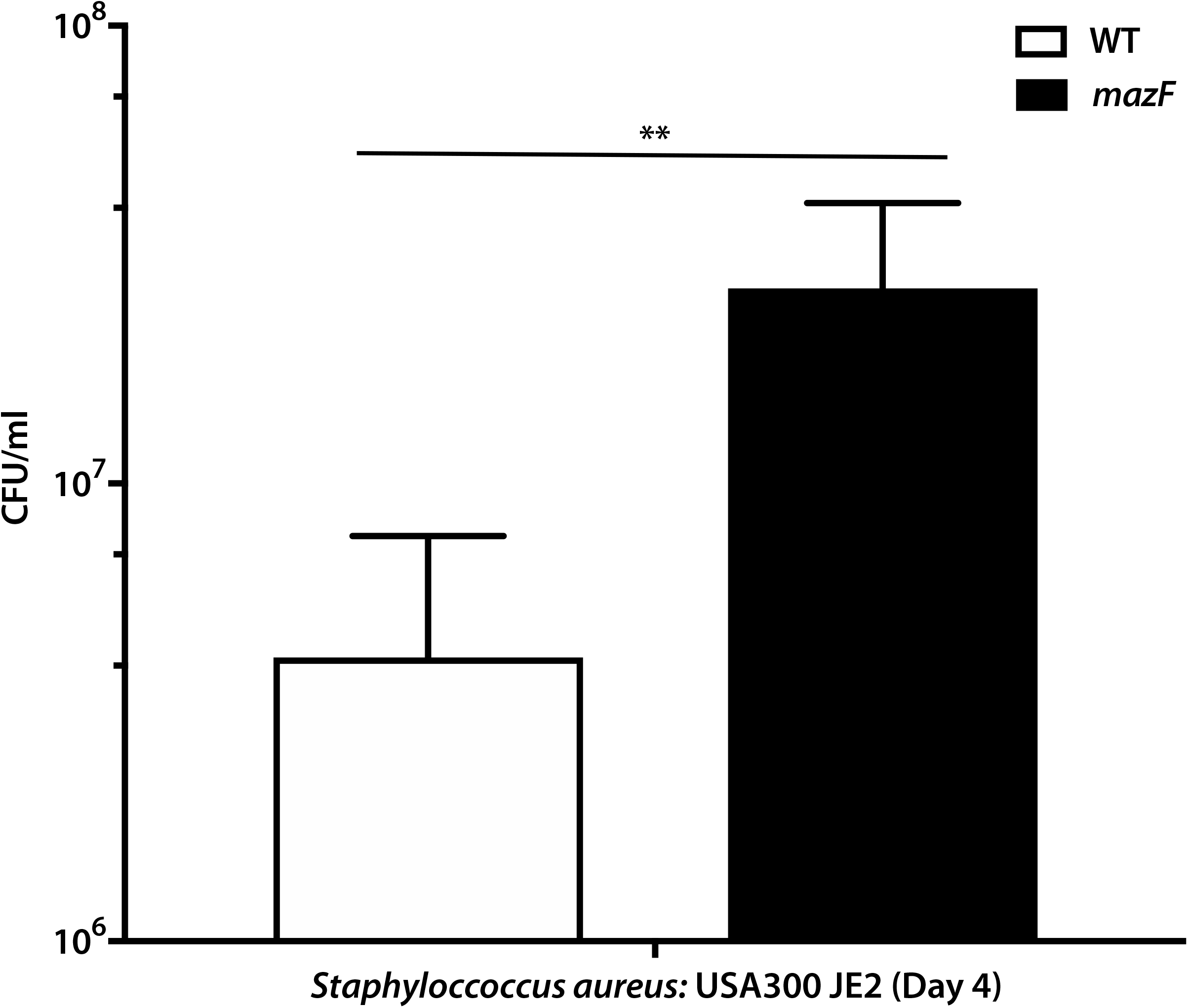
Loss of *mazF* expression increases biofilm formation on surgical implant material in *S. aureus*. Biofilm was cultured on surgical implant material (titanium rods,12mm) for 4 days to form mature biofilm, and the biofilm growth was quantified by sonication, plating, and enumeration for USA 300 JE2. Experiments were completed in triplicate. ** p<0.01. Error bars represent 95% CI (95% confidence interval).

### Loss of *mazF* expression decreases biofilm antibiotic tolerance

In gram negative bacteria, *mazEF* contributes to antibiotic tolerance and bacterial persisters (24–26). The role of *mazEF* and other toxin-antitoxin systems in *S. aureus* antibiotic tolerance is conflicting and unclear (27, 28). We hypothesized that *mazEF* would contribute to biofilm antibiotic tolerance in *S. aureus*. Biofilm antibiotic tolerance was compared between the methicillin resistant *S. aureus* (MRSA) strain JE2 and its corresponding strain disrupted for the *mazF* gene. Mature biofilm cultured on surgical implant material was exposed to 10x minimum inhibitory concentration (MIC) of vancomycin, and quantitative culture was used to assess remaining biofilm mass over three days. Loss of *mazF* expression had a statistically significant increased loss of biofilm mass as compared to the wild type control demonstrating that loss of *mazF* expression decreased biofilm antibiotic tolerance (Fig. 2). These results were confirmed in two additional strains, Newman and SH1000, using both cefazolin and vancomycin (Fig. S4). For all three strains, there was no statistical difference in MICs between the wild type and loss of function strains for cefazolin or vancomycin (Supplemental Table 1).

**Figure 2.**
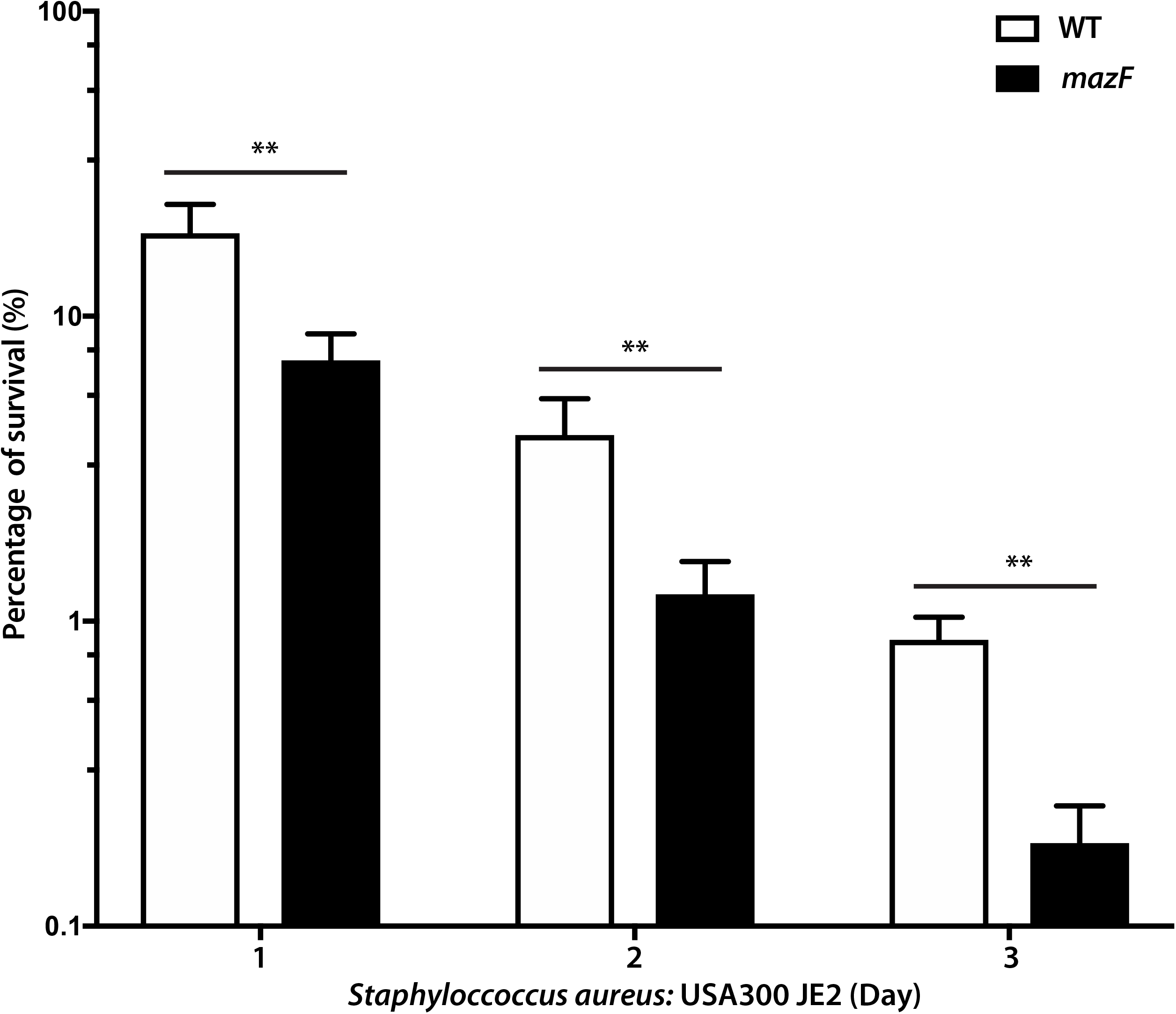
Loss of *mazF* expression decreases biofilm antibiotic tolerance in *S. aureus*. Mature JE2 biofilm was cultured on surgical implant material (4 days on 12mm titanium rods), and exposed to 10x MIC of vancomycin. Implants were then removed, sonicated, and plated to enumerate survivors on a daily basis over 3 days. Remaining biofilm on surgical implant material at each day was compared to the respective pretreated strain. All experiments were completed in triplicate. ** p<0.01. Error bars represent 95% confidence intervals.

### Lack of *mazF* expression only altered planktonic antibiotic tolerance when doubling rate was altered

After observing these strong *mazF* biofilm phenotypes of increased biofilm formation and decreased antibiotic tolerance, we questioned if a similar pattern would be observed in planktonic culture. The growth rate of these three *S. aureus* strains were compared after exiting stationary phase. Loss of *mazF* expression resulted in a statistically significant increased early logarithmic planktonic growth rate in JE2 and SH1000 *S. aureus* strains, but this was not observed at each time point. When the early logarithmic doubling time was compared, only SH1000 and JE2 had a statistically increased doubling rate (Fig. 3A) and Newman did not. A similar pattern was observed with planktonic antibiotic tolerance; deletion or disruption of *mazF* only decreased antibiotic tolerance in the same strains that had an increase in doubling rate, JE2 and SH1000 (Figs. 3B, 3C).

**Figure 3.**
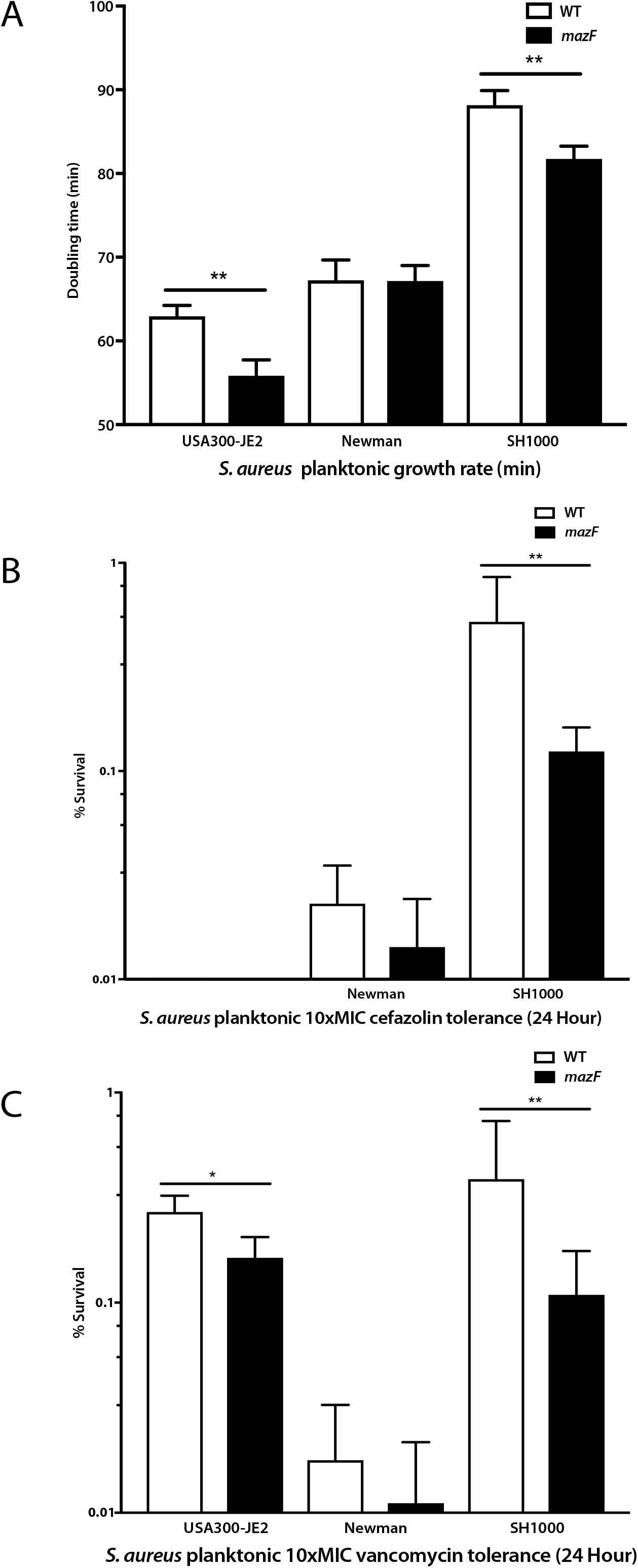
Loss of *mazF* expression increased planktonic growth and decreased vancomycin and cefazolin planktonic antibiotics tolerance in *S. aureus*. (A) Based on the cell growth curve, the doubling time of each strain was determined. Disruption of *mazF* from *S. aureus* resulted in a shorter doubling time in JE2 and SH1000 strains. (B) Disruption of *mazF* expression decreased the cefazolin planktonic antibiotics tolerance in SH1000. The strain JE2 was not included in these experiments as it is methicillin resistant. (C) Disruption of *mazF* expression decreased the planktonic vancomycin tolerance in JE2 and SH1000 strains. All experiments were completed in triplicate. * p<0.05, ** p<0.01. Error bars represent 95% CI (95% confidence interval).

### Loss of *mazF* expression did not alter *sigB* transcription

The *sigB* operon is a master regulator in *S. aureus* that allows it to rapidly redirect transcriptional activities in response to stress (22). It has the potential to be a major regulator of *S. aureus* biofilm formation and virulence (29). The *sigB* operon is directly downstream from *mazEF*, and disruption of *mazF* expression could possibly alter *sigB* expression. We looked at *sigB* expression and genes upstream and downstream of *mazF* to verify that neighboring gene expression was not altered. Quantitative RT-PCR analysis demonstrated no change in expression of *sigB* and the *sigB* dependent gene, asp23 (alkaline shock protein 23), between the three strains with loss of *mazF* expression and their respective wild type control (Fig. 4A and B). We also examined the expression of the genes *rpoF, rsbW*, and *alr* which are directly upstream and downstream of *mazF*, based on genomic location and transcriptional order, using qRT-PCR. There was no statistically significant difference in *rpoF, rsbW* and *alr* expression between these strains and their wild type control (Fig. S5). These results demonstrate that loss of *mazF* did not alter expression of neighboring genes. The observed phenotype was related to the loss of *mazF* and not changes in *sigB* operon or other neighboring gene expression.

**Figure 4.**
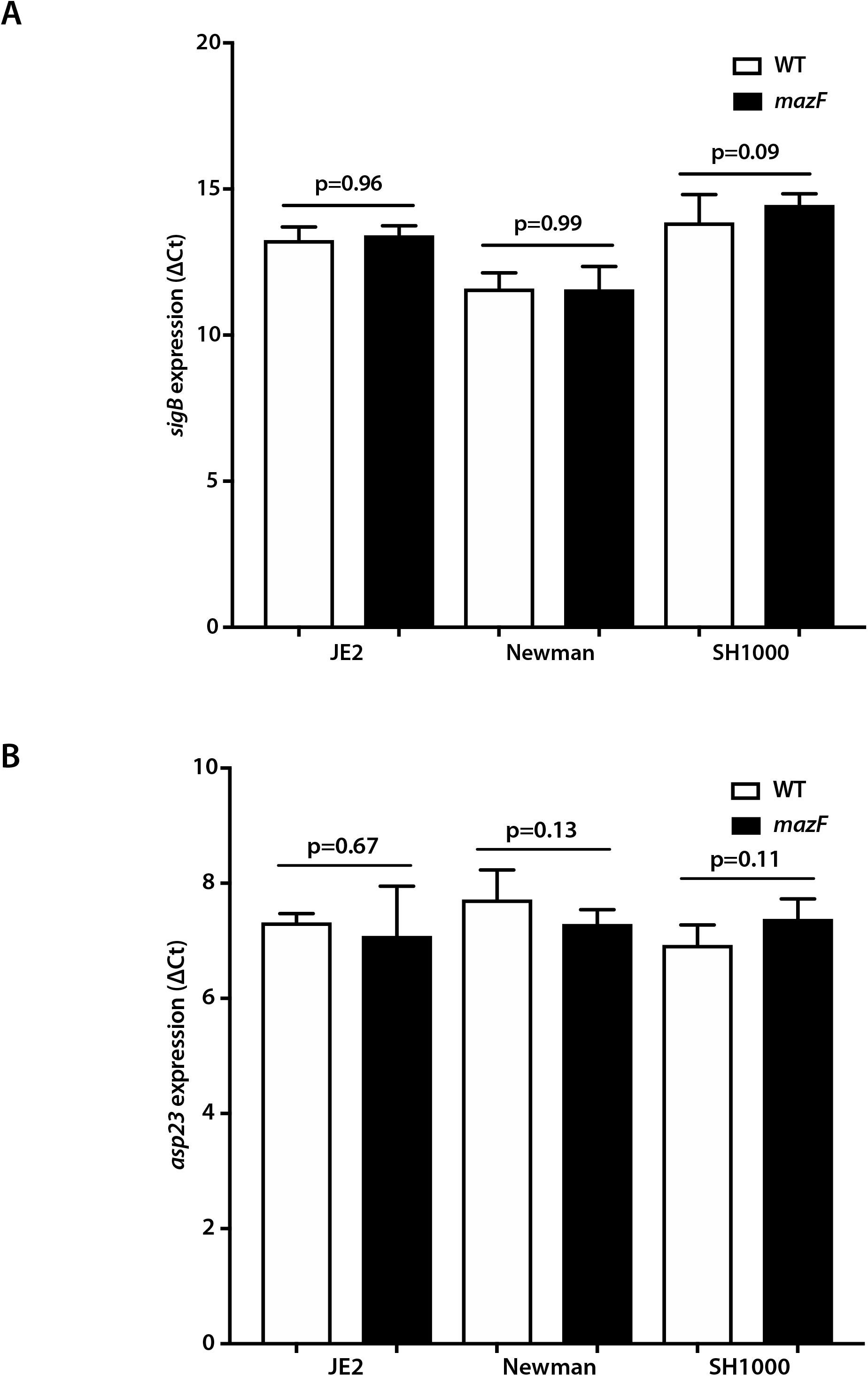
Loss of *mazF* expression had no effect on *sigB* expression. Quantitative real-time RT-PCR analysis of (A) *sigB* and (B) *asp23* expression in three *S. aureus* strains (JE2, Newman, and SH1000). ΔC_t_ value were used to quantify gene expression levels. No significant differences in *sigB* or *asp23* expression were observed between the wild type and loss of *mazF* expression in all three *S. aureus* strains.

### Disruption of *mazF* increased pathogenicity, limited the ability of *S. aureus* to transition from an acute to chronic infection, and inhibited antibiotic tolerance

If lack of *mazF* expression increased biofilm formation, we hypothesized that this increased proliferation would result in increased disease severity. To test this hypothesis, we used a murine abscess model. After inoculation in the hind limb, quantitative culture was used to determine abscess bacterial burden at increasing time points in wild type and the *mazF*::tn strain. We selected the strain JE2 for these experiments as it was the most clinically relevant strain. In immune competent mice, loss of *mazF* had a similar phenotype to *in vitro* observations with increased proliferation and biofilm mass as compared to the wild type strain. After one week, the infection was on a downward trajectory after day 3 (Fig. 5A). To increase disease severity, we repeated experiments in neutropenic mice. Loss of *mazF* expression had increased proliferation and burden as compared to wild-type bacteria. Further, we observed a more virulent and aggressive infection. Wild type mice had almost 100% survival whereas mice inoculated with the *mazF* disrupted strain developed sepsis and death with survival at 25% by day 7 (Fig. 5B). Surprisingly, although a more aggressive infection was observed, the *mazF* disrupted strain was more sensitive to antibiotics than the wild type control. After inoculation, there was a larger decrease in bacterial burden after treatment with vancomycin in the *mazF*::tn *strain* as compared to the wild type (Fig. 5C). Together these results supported the two *in vitro* phenotypes we observed, and suggest that *mazF* contributes to a phenotype of decreased virulence and pathogenesis.

**Figure 5.**
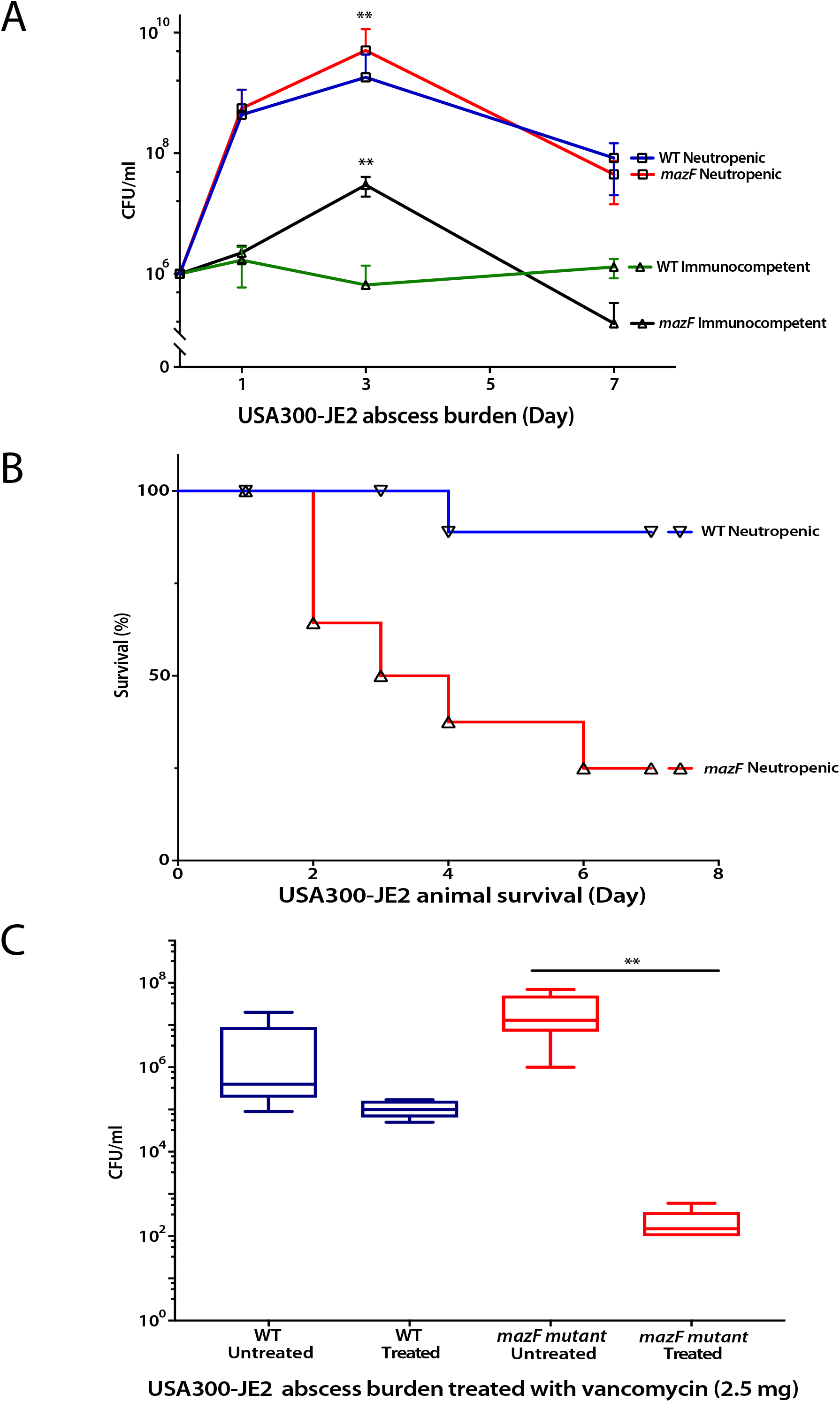
Loss of *mazF* expression increases pathogenicity and limits *S. aureus* ability to establish chronic infection. Bacterial abscess burden and animal survival were used to test the pathogenicity of *S. aureus* wild type and its corresponded *mazF*::tn strains. (A) In both neutropenic and immunocompetent groups, loss of *mazF* increases bacterial burden compared to wild type strains which was most apparent at three days post infection (** P<0.01). (B) Mortality in neutropenic mice inoculated with strains that had no *mazF* expression was 25% on day three and 75% on day seven post infection. Mice inoculated with wild type strain had 0% mortality at day 3 and 10% mortality at day 7. (C) The strain that lost *mazF* expression was more sensitive to antibiotics than the wild type control. After treatment with vancomycin, the loss of *mazF* expression had a 5 log reduction in biofilm as compared to the wild type strain (** P<0.01).

### Increased biofilm formation in a *mazF*-disrupted strain is *ica*-dependent

After a phenotype for *mazF* and a possible role in pathogenesis was identified, we attempted to identify a mechanism behind its regulatory control. The intercellular adhesion gene cluster (*ica*) is composed of *ica*A, *icaD, icaB* and *icaC*, and encodes proteins that promotes intercellular adhesion in many strains and species of *Staphylococcus* (30). Deletion of *mazF* in *S. aureus* results in increased biofilm formation that is *ica*-dependent (31). To test the hypotheses that the phenotype of increased growth and pathogenesis from loss of *mazF* expression was *ica*-dependent, we deleted the *icaADBC* genes from the *mazF*::tn strain to generate a *mazF*::tn/Δ*icaADBC* strain. *S. aureus mazF*::tn/Δ*icaADBC* strains had lower biofilm formation than the wild type and *mazF*::tn strains (Fig. 6A). The neutropenic murine abscess model was repeated with the *mazF*::tn/Δ*icaADBC* strain. The phenotype associated with loss of *mazF* expression was again suppressed. Mice inoculated with the *mazF*::tn/Δ*icaADBC* had comparable survival to the wild type control whereas the *mazF*::tn strain had 50% survival (Fig. 6C). The *mazF*::tn/Δ*icaADBC* strain overcorrected the *mazF*::tn growth phenotype confirming the roles of *icaA, icaB, icaC* and *icaD* in mazEF function and suggests that these 4 genes are likely involved in controlling other process outside the mazEF system. The ability of the *mazF*::tn/Δ*icaADBC* strain to restore survival in the murine abscess model confirmed a role of ica control of biofilm formation in pathogenesis.

**Figure 6.**
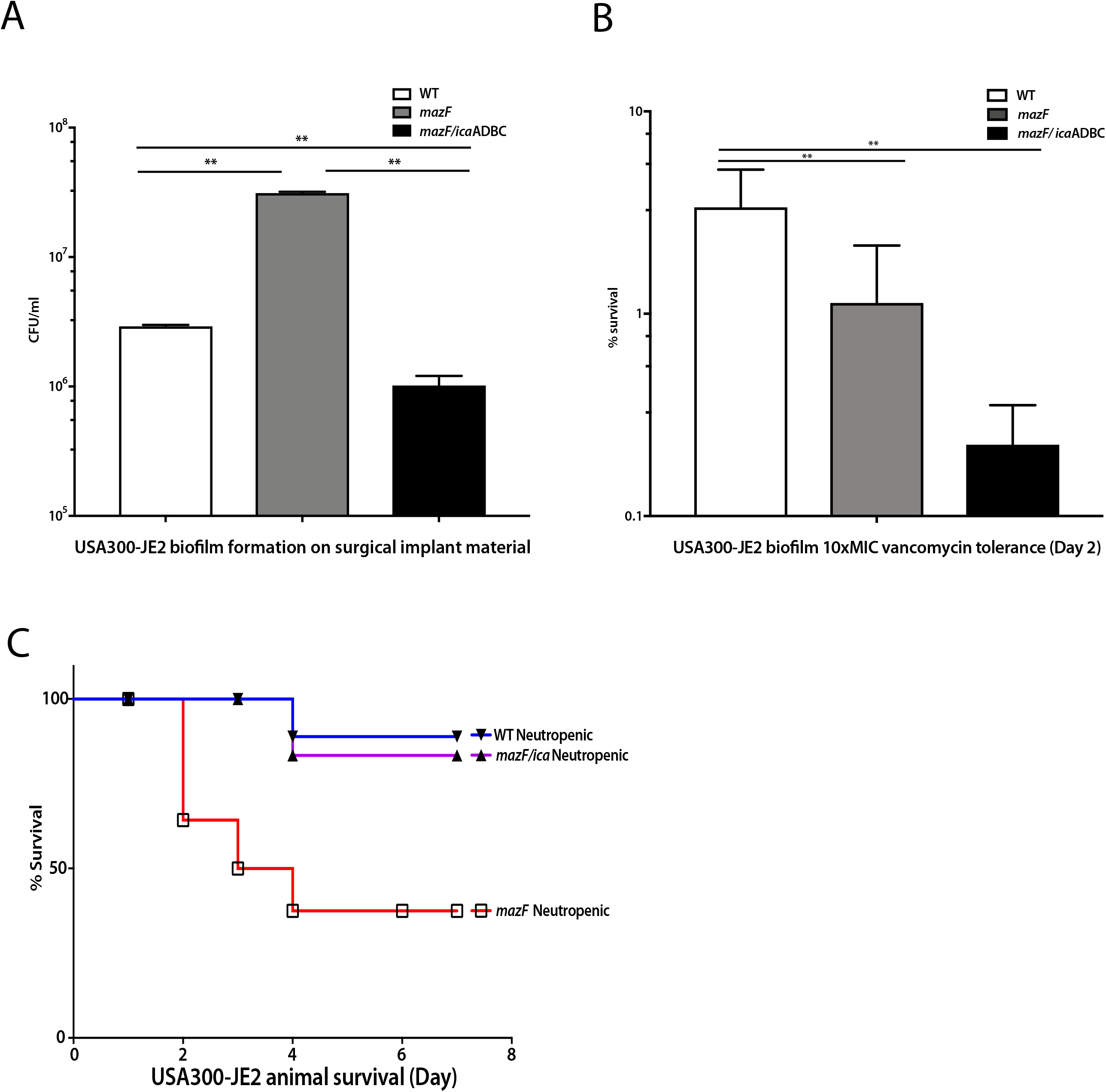
Increased biofilm formation in the *mazF* disruption strain is *ica*-dependent. Biofilm formation in the *mazF*::tn and *mazF*::tn/Δ*icaADBC* strain were compared to the parental strain. (A) Biofilm formation of the *mazF*::tn/Δ*icaADBC* strain was lower than that of strains lacking *mazF* expression alone. (B) Biofilm antibiotic tolerance of the double loss of function was lower than that of strains lacking *mazF* expression alone. (C). *mazF/icaADBC* double loss of function strain reversed the animal mortality to wild type strain levels. All experiments were completed in triplicate. ** p<0.01. Error bars represent 95% CI (95% confidence interval).

### Decreased biofilm antibiotic tolerance in a *mazF* disruption is not *ica*-dependent

We then tested the role of *icaADBC* in regulating *mazF* biofilm antibiotic tolerance. Mature biofilm was exposed to 10x MIC vancomycin. *mazF*::tn and *mazF*::tn/Δ*icaADBC* strains had decreased vancomycin tolerance compared with wild type strain. Unlike the biofilm formation phenotype where deletion of *icaADBC* reversed the *mazF* disruption phenotype, loss of both *mazF* and *icaADBC* expression resulted in even less biofilm antibiotic tolerance than loss of *mazF* alone. This demonstrated that ica genes were also involved in antibiotic tolerance as well (Fig. 6B).

## Discussion

The physiologic role of bacterial toxin-antitoxin systems remain unknown. In *S. aureus, mazEF* is a well-studied toxin-antitoxin system whose phenotype and physiologic role in *S. aureus* remains elusive. Loss of *mazF* expression resulted in a phenotype of increased biofilm formation on surgical implant material and decreased biofilm antibiotic tolerance in all three *S. aureus* strains, but little change in planktonic cells. In our murine abscess model, the phenotypes associated with *mazEF* contributed to a biofilm-dependent disease process that is consistent with most chronic bacterial infections and the clinical manifestation of surgical infections. Further mechanistic analysis supported a role for extracellular polysaccharide adhesins in the increased biofilm formation and pathogenesis when *mazF* expression is disrupted. Combined, these results suggest that *mazEF* helps regulate the transition between acute to chronic infection in *S. aureus*.

Regulation of growth and biofilm formation is a phenotype associated with TA systems. The mechanism of toxin-antitoxin systems includes an antitoxin that prevents the toxin from inducing growth arrest using a variety of tools (24, 32). After loss of *mazF* expression, we observed increased biofilm when compared to its isogenic wild-type strain on fibrinogen-coated plastic and titanium *in vitro* and *in vivo*. This supports the work of other groups where overexpression of *mazF* in *S. aureus* resulted in growth arrest (21) and *mazF* mutants had increased biofilm formation (31). This is the primary mechanism where bacteria are thought to become antibiotic tolerant from toxin-antitoxin systems; dormant bacteria are tolerant to an antibiotic whose main process is disrupting their metabolism.

There is evidence to suggest that TA systems play an important role in antibiotic tolerance based on multiple examples in gram negative species (26) as well as in acid-fast mycobacterium (17). Although there is less evidence for this phenotype in gram-positive organisms, it has been suspected a similar pattern exists in *S. aureus*. We observed a difference in biofilm antibiotic tolerance when *mazF* expression was disrupted as compared to its wild type strains (Fig. 2). We did not observe a difference in the MIC. This supports similar previous observations (31). Other groups have noted that in *S. aureus mazF* transcription is altered by sub-MIC concentrations of MICs of tetracycline, penicillin, and linezolid (22). Despite generating greater biofilm formation, the loss of *mazF* expression demonstrated increased antibiotic susceptibility to clinically relevant antibiotics cefazolin and vancomycin. Combined this provides strong evidence for a major role of *mazF* in *S. aureus* biofilm antibiotic tolerance.

Persisters are a subpopulation of bacteria that have a phenotypic tolerance to antibiotics (33, 34). The mechanism behind this tolerance is thought to be regulated by the metabolic state of the cell (35). We observed that the role of *mazF* in antibiotic tolerance appears to be correlated with growth. The phenotype of antibiotic tolerance was more weakly observed in planktonic *S. aureus* strains (Fig. 3). In planktonic culture, only loss of function strains with decreased doubling times as compared to the wild type strains were observed to have decreased planktonic antibiotic tolerance. This supports other results suggesting that persister formation is based on ATP levels, and is an area of future work. Likely, multiple mechanisms exist to support persister cell formation and antibiotic tolerance, including the stringent response (28).

Biofilm formation is an important step for *S. aureus* to establish an infection. This is regulated by polysaccharide intercellular adhesin (PIA/PNAG) encoded by the *ica* operon (36). Based on this and our observation that loss of *mazF* expression increased biofilm formation, we speculated that *mazF* inhibits biofilm formation by decreasing *ica* transcription. A *mazF*::tn/Δ*icaADBC* strain reversed the *in vitro* and *in vivo* phenotypes from loss of *mazF* expression (Fig. 6). This *mazF*::tn Δ*icaADBC* strain had similar pathogenicity as the wild type strain. This provides evidence that *ica*-mediated biofilm formation and pathogenicity are inhibited by *mazF*. This supports other groups observations that *S. aureus* biofilm formation is dependent on *mazF* mRNA interferase activity (31).

*S. aureus* infections are typically acute. Although there is a range of pathogenesis from simple, superficial abscesses to life threatening systemic sepsis, the outcomes of these disease processes resolve over a limited time period. An exception is surgical infection where chronic infections can develop over an extended period of time and biofilm formation plays an important physiologic role (5, 11). Regulation of growth, biofilm formation, and antibiotic tolerance could have important roles of bacteria physiology in this disease state. *S. aureus* biofilm formation is an essential step in establishing infection and pathogenicity (37, 38). Surprisingly, although loss of *mazF* created a more virulent organism with higher lethality, these infections were also more susceptible to antibiotics. Combined, these results suggest that *mazF* expression inhibits biofilm formation and increases antibiotic tolerance allowing the bacteria to transition to a chronic infection that is more challenging to treat. This demonstrates a physiologic role for toxin antitoxin systems during infections. *MazEF* toxin antitoxin systems not only make the bacteria more tolerant to antibiotics but makes the bacteria more tolerant to the host.

## Materials and Methods

### Bacterial strains, plasmids and growth conditions

USA 300 JE2 was selected as the primary strain as it was the most clinically relevant, and USA300 clones have the highest growth rate as compared to other common *S. aureus* strains (39). All bacterial strains and plasmids used in this study are listed in Table 1. *Staphylococcus aureus* strains were cultured in Trypticase soy broth (TSB) medium with or without antibiotics.

**Table 1.**
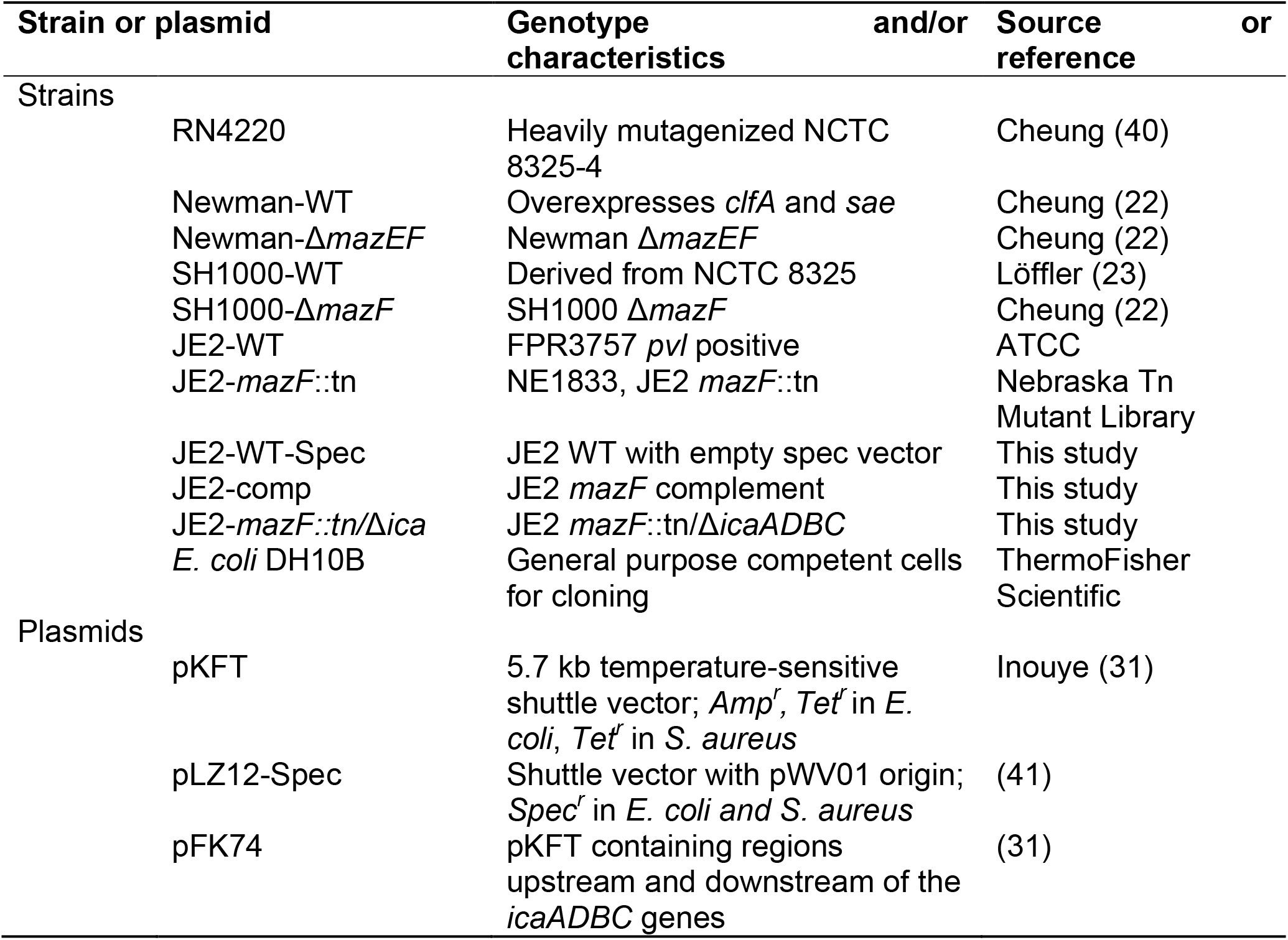
Bacterial stains and plasmids used in this study

### Genomic bacterial DNA isolation

Genomic DNA was isolated from *S. aureus* samples by following manufacturer’s instructions (MasterPure gram positive DNA Purification Kit; Lucigen, USA). Briefly, a single colony from a TSB plate was inoculated in TSB medium and grown overnight at 37°C in an orbital shaker. Pellet 1.5 ml culture and resuspend in 150 μl TE buffer. Lysis the bacteria in lysis buffer at 37°C until bacterial cell wall is destroyed. Treated with Proteinase K. After added protein precipitation reagent. Pellet the debris by centrifugation at 4°C for 10 minutes at 12,000 x g. Keep the supernatant and pellet the genome DNA with isopropanol. Rinse the pellet with 70% ethanol. Resuspend the DNA in TE Buffer. The genomic DNA can be used as template in following PCR reactions.

### Creation the *mazF* complementary strain

The complete *mazF* gene was amplified from 860 bp upstream of the *mazF* open reading frame, including promotor region, in JE2 by PCR and cloned into a pLZ12-spec shuttle vector. The transformed *mazF* expression vector was transformed into the *mazF*::tn strain and selected with 200 μg/ml spectinomycin.

### Isolation of RNA and quantitative RT-PCR analysis

RNA isolation and quantitative reverse transcription polymerase chain reaction (RT-PCR) were performed by following the manufacturer’s instruction of the product. In brief, *S. aureus* was grown in 4 mL of TSB medium supplemented with appropriate antibiotics at 37°C for 16 hours. Overnight culture was centrifuged, and pellet resuspended in TE Buffer by vortexing. 500 μg/ml lysostaphin (Sigma-Aldrich) was added to the resuspended bacteria and incubate at 37°C for 15 minutes. Total RNA was extracted using TRIzol^®^ Max™ Bacterial RNA Isolation Kit (Thermo Fisher Scientific). Single-stranded cDNA was created from reverse transcription of the RNA using SuperScript IV Reverse Transcriptase (Thermo Fisher Scientific). The newly synthesized cDNA was used immediately or frozen at −80 °C.

Quantitative RT-PCR analysis was performed using the CFX96 Real-Time System (BioRad, Richmond, CA) and PowerUp™ SYBR Green Master Mix (Thermo Fisher Scientific). The cycling conditions were 50 °C for 10 min and 95 °C for 5 min, followed by 45 cycles of 95 °C for 10 sec and 62 °C for 30 sec. For all samples, the threshold cycle number (C_t_) at which the fluorescence values became logarithmic was determined. The ΔC_t_ value was calculated for each sample as the difference between the sample C_t_ and the control C_t_.

### Creation of *icaADBC* gene deletion using pKFT Vector

A JE2 double *mazF*::tn/Δ*icaADBC* strain was created from the base JE2 *mazF*::tn strain using a previously described protocol (42). Briefly, the allelic replacement vector pFK74 containing regions upstream 1.1 kb and downstream 0.9 kb of the *icaADBC* gene was first transformed into DNA restriction system-deficient *S. aureus* RN4220, then a modified plasmid was isolated and electroporated into *mazF*::tn strain (NE1833) from NARSA NR-48501 library. Transformants were selected at 30 °C on TSB plates containing tetracycline. A single colony transformant was cultured at 30 °C in TSB media containing tetracycline in an orbital shaker. Integration of the plasmid into the chromosome by a single crossover event was achieved by incubation at 42 °C on TSB plates containing tetracycline. Correct homologous recombination of the target region was verified by PCR using primer set of pUC-UV (5’-CGACGTTGTAAAACGACGGCCAGT-3’, plasmid) and *icaADBC* 5’-up (5’-CCATCACATAGGCGCTTATCAA-3’, chromosome) or pUC-RV (5’-CATGGTCATAGCTGTTTCCTGTG-3’, plasmid) and *icaADBC* 3’-dn (5’-GAAGCAACGCACAAAGCATTA-3’, chromosome). Integrants were grown at 25 °C overnight with shaking in 10 ml TSB without any antibiotics. Bacteria was serially diluted and plated on TSB plates at 42 °C. The excision of the plasmid region in the chromosome by a second crossover event was screened for by isolation of tetracycline-sensitive colonies by replica-plating candidates on TSB plates versus TSB plates containing tetracycline (3 μg/ml). Integrants were cultured overnight at 37 °C. Then, the markerless deletion mutants were screened by PCR using primers *icaADBC-1* (5’-AAAAAGATCTTTAGTAGCGAATACACTTC-3’) and *icaADBC-4* (5’-TACAAGATCTTTGGCATCATTTAGCAGAC-3’) from tetracycline-sensitive colonies. The strain with *icaADBC* deletion was screened by PCR and confirmed by DNA sequencing.

### Cell growth curve and doubling time

Approximately 1×10^6^ cells were added from overnight culture to fresh TSB medium and incubated at 37°C. The OD_600_ absorbance (TECAN, infinite M200) was measured every hour during a 24 hour period. Calculation of the doubling time was based off of these measurements.

### Biofilm formation assay

Four titanium rods (12 mm) per well were incubated in TSB growth medium inoculated with 1×10^4^ CFU *S. aureus* for 24 to 96 hours. Titanium rods were then washed three times with 1ml PBS and then sonicated for 30 minutes in 1 ml fresh TSB medium. After serial 1:10 dilution, the bacterial concentration (CFU/mL) was determined via colony-forming unit (CFU) assay on TSA II blood agar plates (Thermo Fisher Scientific, USA). A semi-quantitative adherence assay was performed on 96-well tissue culture polystyrene plates (Sigma-Aldrich, USA). Plates were coated with 200 μl of phosphate-buffered saline (PBS) containing 5 μg/ml fibrinogen (Sigma-Aldrich, USA) overnight at 4°C. Washed three times with PBS and then blocked with 100 μl of a 2% bovine serum albumin (BSA) solution for 1 h at 37°C. The wells were carefully washed three times with 100 μl of PBS; 100 μl of bacteria (approximately 1×10^7^ cells) was added to the appropriate wells and incubated for 24 hours at 37 °C. The wells were washed four times with 100 μl of PBS. Bacteria were fixed with 100 μl of 10 *%* formaldehyde (Sigma-Aldrich, USA) for 10 min. Then 100 μl of 0.2% crystal violet (Sigma-Aldrich, USA) was added to each well for 10 min, cells were washed four times with distilled water. Air dried wells for 2 hours and then add 100 μl of 30% acetic acid (Fisher Scientific, USA) to dissolve crystal violet and the absorbance was measured at 590 nm.

### Minimum inhibitory concentration (MIC) assay

A single colony from an overnight agar plate was inoculated in 5 ml TSB medium to achieve the specified inoculum turbidity by comparing to a 0.5x McFarland turbidity standard (~1×10^8^ CFU/ml). A sterile swab was placed in the inoculum suspension and streaked across the entire agar surface six times, rotating the plate to evenly distribute the inoculum. An Etest MIC test strips (Liofilchem, Italy) was applied with sterile forceps. Agar plates were then incubated in an inverted position at 37 °C overnight.

### Biofilm and planktonic antibiotics tolerance assay

For biofilm assay, grow *S. aureus* strains on surgical implant material (12mm titanium rods) for 4 days to form mature biofilm (change medium every day). Then, exposed the mature biofilm to 10x MIC of cefazolin or vancomycin for 3 days (change medium every day). Implants were then washed, removed, sonicated, and plated to enumerate survivors (CFU assay) at each day. For planktonic assay, grow *S. aureus* strains in fresh TSB media overnight. Next day, 1:100 diluted the overnight culture media with fresh TSB medium and grow to around 0.5X McFarland turbidity under 37°C. Do CFU assay to determine the viable cells before the treatment. Then, exposed the bacteria to 10X MIC cefazolin or vancomycin to 4-hour and 24-hour time point. And plated to enumerate survivors (CFU assay) at each time point. Calculate the percentage of survival (%) at each time point. All experiments were completed in triplicate. * p<0.05, ** p<0.01. Error bars represent 95% CI (95% confidence interval).

### Mice, Neutropenic Thigh model and *Staphylococcus aureus* strains administration

Eight-week-old B57BL/6J mice were purchased from the Jackson laboratory (Bar Harbor, ME, USA). All animal protocols used for these experiments were approved by the University of Pittsburgh’s Institutional Animal Care and Use Committee. Mice were rendered neutropenic by two 100 μl intra peritoneal injections of cyclophosphamide (150 mg/kg three days pre-infection and 100 mg/kg one day pre-infection). Mice where anesthetized by 2% isoflurane, hair was removed from leg and treated with betadine. An inoculation volume of 100 μl, 1×10^6^ CFU of JE2-WT or JE2-Δ*mazF*::tn strain was injected into the thigh. Mice were monitored for weight loss, leg swelling, ambulatory abilities, signs of sepsis, and death. Mice were sacrificed at 1, 3, and 7 days post infection. A ~5 x 5 mm piece of thigh muscle from infection site was obtained and placed into 1% Tween 20 in PBS on ice. Abscess samples were sonicated 10 minutes, and colony forming unit (CFU) assay was performed on blood agar plates to quantify bacterial burden.

### Statistical analysis

Statistical analysis was based on the number of populations and comparisons. Student t-test was sued for two populations. One-way anova and two-way anova was used for comparing multiple populations across either one condition or two conditions respectively. To determine antibiotic tolerance, multilevel mixed-effects linear regression models were constructed to compare the rate of change in CFU/mL over time between wild type and strains with loss of *mazF* expression. The outcome, CFU/mL, was natural-log transformed to produce approximately normally distributed values before fitting models via maximum likelihood estimation (MLE). Bacteria type (WT, Δ*mazF*), time, and a type-by-time interaction were included as fixed effects in the multilevel models, and the baseline concentration was accounted for. Random effects for Experiment, as well as Group (nested within Experiment), were included in all models to adjust for within-cluster correlation. Of primary interest were the type-by-time interaction coefficients, which reflect the degree to which the rate of decline in log-CFU differs between wildtype and Δ*mazF* bacteria. After fitting the models, the estimated interaction coefficients were back-transformed to provide interpretable results on the original, non-logarithmic CFU/mL scale.

## Acknowledgements

Dr. Kenneth Urish is supported in part by the National Institute of Arthritis and Musculoskeletal and Skin Diseases (NIAMS K08AR071494), the National Center for Advancing Translational Science (NCATS KL2TR0001856), and the Orthopaedic Research and Education Foundation.

## Supporting information

**Supplemental Figure 1. Loss of *mazF* increases biofilm formation on surgical implant material in *S. aureus*.**

Biofilm was cultured on surgical implant material (titanium rods,12 mm) for 4 days to form mature biofilm, and the biofilm growth was quantified by sonication, plating, and enumeration for JE2, Newman and SH1000 strains, respectively. All experiments were completed in triplicate. ** p<0.01. Error bars represent 95% CI (95% confidence interval).

**Supplemental Figure 2. Bacterial biofilm formation on 96-well culture plate**

*S. aureus* strains were cultured on fibrinogen coated 96-well polystyrene plates for 24 to 48 hours. Biofilm formation was quantified using the crystal violet method, the absorbance was measured at 590 nm. All experiments were completed in triplicate. ** p<0.01. Error bars represent 95% CI (95% confidence interval).

**Supplemental Figure 3. MazF complement reduces the planktonic cell growth and biofilm formation**

A genomic complement approach was used to restore *mazF* expression in JE2, and the growth phenotype was reversed. JE2 wild type strain with empty spec vector as a control. Biofilm formation measured by using the crystal violet assay (A) and titanium rods CFU assay (B) and planktonic growth measured using optical density (C) demonstrated that biofilm formation and planktonic growth was decreased in the *mazF* complement strain. All experiments were completed in triplicate. * p<0.05, ** p<0.01. Error bars represent 95% CI (95% confidence interval).

**Supplemental Figure 4. Loss of *mazF* decreases biofilm vancomycin and cefazolin tolerance in *S. aureus*.**

Mature biofilm grown on surgical implant material (4 days on titanium rods) was exposed to 10x MIC of cefazolin or vancomycin. Implants were then removed, sonicated, and plated to enumerate survivors on a daily basis over 3 days. A regression model was used to estimate the overall percent of biofilm remaining at day 3 relative to pretreated. (A) Biofilm of Newman and SH1000 strains were exposed to cefazolin. JE2 was not included as it is a MRSA. (B) Biofilm of JE2, Newman, and SH1000 strains were exposed to vancomycin. All experiments were completed in triplicate. * p<0.05, ** p<0.01. Error bars represent 95% confidence intervals.

**Supplemental Figure 5. Deletion of *mazF* had no effect on *sigB* operon expression**

Quantitative real-time RT-PCR analysis of the *sigB* operon with *rpoF, rsbW*, and *alr* transcripts. ΔC_t_ value was used to indicate the expression levels of selected genes.

**Supplemental Table 1.**
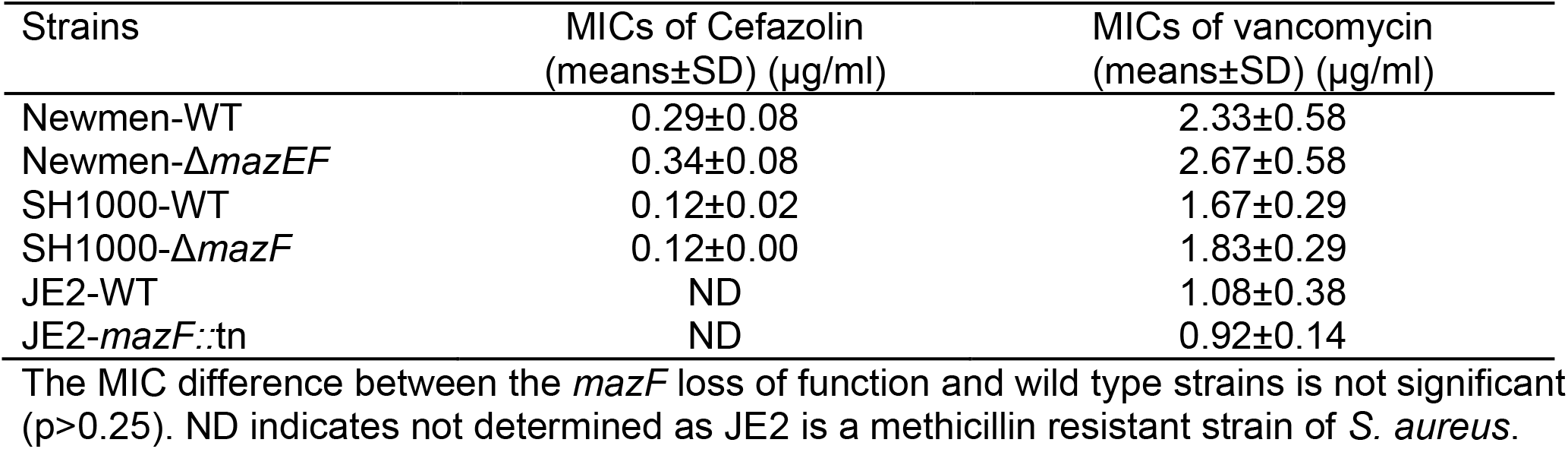
Cefazolin and vancomycin MICs of non-biofilm *S. aureus*

| Strains | MICs of Cefazolin<br>(means±SD) (μg/ml) | MICs of vancomycin<br>(means±SD) (μg/ml) |
| --- | --- | --- |
| Newmen-WT | 0.29±0.08 | 2.33±0.58 |
| Newmen-$\Delta mazEF$ | 0.34±0.08 | 2.67±0.58 |
| SH1000-WT | 0.12±0.02 | 1.67±0.29 |
| SH1000-$\Delta mazF$ | 0.12±0.00 | 1.83±0.29 |
| JE2-WT | ND | 1.08±0.38 |
| JE2-*mazF*::tn | ND | 0.92±0.14 |

The MIC difference between the *mazF* loss of function and wild type strains is not significant ( $p > 0.25$ ). ND indicates not determined as JE2 is a methicillin resistant strain of *S. aureus*.

## References

1. Lowy FD. 1998. Staphylococcus aureus infections. N Engl J Med 339:520–532.

2. Magill SS, O’Leary E, Janelle SJ, Thompson DL, Dumyati G, Nadle J, Wilson LE, Kainer MA, Lynfield R, Greissman S, Ray SM, Beldavs Z, Gross C, Bamberg W, Sievers M, Concannon C, Buhr N, Warnke L, Maloney M, Ocampo V, Brooks J, Oyewumi T, Sharmin S, Richards K, Rainbow J, Samper M, Hancock EB, Leaptrot D, Scalise E, Badrun F, Phelps R, Edwards JR, Emerging Infections Program Hospital Prevalence Survey T. 2018. Changes in Prevalence of Health Care-Associated Infections in U.S. Hospitals. N Engl J Med 379:1732–1744.

3. Bozic KJ, Kurtz SM, Lau E, Ong K, Chiu V, Vail TP, Rubash HE, Berry DJ. 2010. The epidemiology of revision total knee arthroplasty in the United States. Clin Orthop Relat Res 468:45–51.

4. Koh CK, Zeng I, Ravi S, Zhu M, Vince KG, Young SW. 2017. Periprosthetic Joint Infection Is the Main Cause of Failure for Modern Knee Arthroplasty: An Analysis of 11,134 Knees. Clin Orthop Relat Res 475:2194–2201.

5. Urish KL, Bullock AG, Kreger AM, Shah NB, Jeong K, Rothenberger SD, Infected Implant C. 2018. A Multicenter Study of Irrigation and Debridement in Total Knee Arthroplasty Periprosthetic Joint Infection: Treatment Failure Is High. J Arthroplasty 33:1154–1159.

6. Yoon HK, Cho SH, Lee DY, Kang BH, Lee SH, Moon DG, Kim DH, Nam DC, Hwang SC. 2017. A Review of the Literature on Culture-Negative Periprosthetic Joint Infection: Epidemiology, Diagnosis and Treatment. Knee Surg Relat Res 29:155–164.

7. Zmistowski B, Karam JA, Durinka JB, Casper DS, Parvizi J. 2013. Periprosthetic joint infection increases the risk of one-year mortality. J Bone Joint Surg Am 95:2177–2184.

8. Choi HR, Bedair H. 2014. Mortality following revision total knee arthroplasty: a matched cohort study of septic versus aseptic revisions. J Arthroplasty 29:1216–1218.

9. Lum ZC, Natsuhara KM, Shelton TJ, Giordani M, Pereira GC, Meehan JP. 2018. Mortality During Total Knee Periprosthetic Joint Infection. J Arthroplasty 33:3783–3788.

10. Kurtz SM, Lau EC, Son MS, Chang ET, Zimmerli W, Parvizi J. 2018. Are We Winning or Losing the Battle With Periprosthetic Joint Infection: Trends in Periprosthetic Joint Infection and Mortality Risk for the Medicare Population. J Arthroplasty 33:3238–3245.

11. Ma D, Shanks RMQ, Davis CM, 3rd, Craft DW, Wood TK, Hamlin BR, Urish KL. 2018. Viable bacteria persist on antibiotic spacers following two-stage revision for periprosthetic joint infection. J Orthop Res 36:452–458.

12. Urish KL, DeMuth PW, Kwan BW, Craft DW, Ma D, Haider H, Tuan RS, Wood TK, Davis CM, 3rd. 2016. Antibiotic-tolerant Staphylococcus aureus Biofilm Persists on Arthroplasty Materials. Clin Orthop Relat Res 474:1649–1656.

13. Van Melderen L, Saavedra De Bast M. 2009. Bacterial toxin-antitoxin systems: more than selfish entities? PLoS Genet 5:e1000437.

14. Fozo EM, Makarova KS, Shabalina SA, Yutin N, Koonin EV, Storz G. 2010. Abundance of type I toxin-antitoxin systems in bacteria: searches for new candidates and discovery of novel families. Nucleic Acids Res 38:3743–3759.

15. Gerdes K. 2000. Toxin-antitoxin modules may regulate synthesis of macromolecules during nutritional stress. J Bacteriol 182:561–572.

16. Zhang Y, Zhang J, Hoeflich KP, Ikura M, Qing G, Inouye M. 2003. MazF cleaves cellular mRNAs specifically at ACA to block protein synthesis in Escherichia coli. Mol Cell 12:913–923.

17. Tiwari P, Arora G, Singh M, Kidwai S, Narayan OP, Singh R. 2015. MazF ribonucleases promote Mycobacterium tuberculosis drug tolerance and virulence in guinea pigs. Nat Commun 6:6059.

18. Tiwari P, Arora G, Singh M, Kidwai S, Narayan OP, Singh R. 2015. Corrigendum: MazF ribonucleases promote Mycobacterium tuberculosis drug tolerance and virulence in guinea pigs. Nat Commun 6:7273.

19. Gerdes K, Christensen SK, Lobner-Olesen A. 2005. Prokaryotic toxin-antitoxin stress response loci. Nat Rev Microbiol 3:371–382.

20. Korch SB, Malhotra V, Contreras H, Clark-Curtiss JE. 2015. The Mycobacterium tuberculosis relBE toxin:antitoxin genes are stress-responsive modules that regulate growth through translation inhibition. J Microbiol 53:783–795.

21. Fu Z, Tamber S, Memmi G, Donegan NP, Cheung AL. 2009. Overexpression of MazFsa in Staphylococcus aureus induces bacteriostasis by selectively targeting mRNAs for cleavage. J Bacteriol 191:2051–2059.

22. Donegan NP, Cheung AL. 2009. Regulation of the mazEF toxin-antitoxin module in Staphylococcus aureus and its impact on sigB expression. J Bacteriol 191:2795–2805.

23. Tuchscherr L, Medina E, Hussain M, Volker W, Heitmann V, Niemann S, Holzinger D, Roth J, Proctor RA, Becker K, Peters G, Loffler B. 2011. Staphylococcus aureus phenotype switching: an effective bacterial strategy to escape host immune response and establish a chronic infection. EMBO Mol Med 3:129–141.

24. Wang X, Wood TK. 2011. Toxin-antitoxin systems influence biofilm and persister cell formation and the general stress response. Appl Environ Microbiol 77:5577–5583.

25. Lewis K. 2008. Multidrug tolerance of biofilms and persister cells. Curr Top Microbiol Immunol 322:107–131.

26. Tripathi A, Dewan PC, Siddique SA, Varadarajan R. 2014. MazF-induced growth inhibition and persister generation in Escherichia coli. J Biol Chem 289:4191–4205.

27. Schuster CF, Mechler L, Nolle N, Krismer B, Zelder ME, Gotz F, Bertram R. 2015. The MazEF Toxin-Antitoxin System Alters the beta-Lactam Susceptibility of Staphylococcus aureus. PLoS One 10:e0126118.

28. Conlon BP, Rowe SE, Gandt AB, Nuxoll AS, Donegan NP, Zalis EA, Clair G, Adkins JN, Cheung AL, Lewis K. 2016. Persister formation in Staphylococcus aureus is associated with ATP depletion. Nat Microbiol 1:16051.

29. Mitchell G, Fugere A, Pepin Gaudreau K, Brouillette E, Frost EH, Cantin AM, Malouin F. 2013. SigB is a dominant regulator of virulence in Staphylococcus aureus small-colony variants. PLoS One 8:e65018.

30. Cramton SE, Gerke C, Schnell NF, Nichols WW, Gotz F. 1999. The intercellular adhesion (ica) locus is present in Staphylococcus aureus and is required for biofilm formation. Infect Immun 67:5427–5433.

31. Kato F, Yabuno Y, Yamaguchi Y, Sugai M, Inouye M. 2017. Deletion of mazF increases Staphylococcus aureus biofilm formation in an ica-dependent manner. Pathog Dis 75.

32. Page R, Peti W. 2016. Toxin-antitoxin systems in bacterial growth arrest and persistence. Nat Chem Biol 12:208–214.

33. Brauner A, Fridman O, Gefen O, Balaban NQ. 2016. Distinguishing between resistance, tolerance and persistence to antibiotic treatment. Nat Rev Microbiol 14:320–330.

34. Balaban NQ, Merrin J, Chait R, Kowalik L, Leibler S. 2004. Bacterial persistence as a phenotypic switch. Science 305:1622–1625.

35. Zampieri M, Enke T, Chubukov V, Ricci V, Piddock L, Sauer U. 2017. Metabolic constraints on the evolution of antibiotic resistance. Mol Syst Biol 13:917.

36. Fluckiger U, Ulrich M, Steinhuber A, Doring G, Mack D, Landmann R, Goerke C, Wolz C. 2005. Biofilm formation, icaADBC transcription, and polysaccharide intercellular adhesin synthesis by staphylococci in a device-related infection model. Infect Immun 73:1811–1819.

37. Costerton JW. 1999. Introduction to biofilm. Int J Antimicrob Agents 11:217–221; discussion 237–219.

38. Donlan RM. 2001. Biofilm formation: a clinically relevant microbiological process. Clin Infect Dis 33:1387–1392.

39. Thurlow LR, Joshi GS, Richardson AR. 2012. Virulence strategies of the dominant USA300 lineage of community-associated methicillin-resistant Staphylococcus aureus (CA-MRSA). FEMS Immunol Med Microbiol 65:5–22.

40. Nair D, Memmi G, Hernandez D, Bard J, Beaume M, Gill S, Francois P, Cheung AL. 2011. Whole-genome sequencing of Staphylococcus aureus strain RN4220, a key laboratory strain used in virulence research, identifies mutations that affect not only virulence factors but also the fitness of the strain. J Bacteriol 193:2332–2335.

41. Husmann LK, Scott JR, Lindahl G, Stenberg L. 1995. Expression of the Arp protein, a member of the M protein family, is not sufficient to inhibit phagocytosis of Streptococcus pyogenes. Infect Immun 63:345–348.

42. Kato F, Sugai M. 2011. A simple method of markerless gene deletion in Staphylococcus aureus. J Microbiol Methods 87:76–81.

